# FunFun: ITS-based functional annotator of fungal communities

**DOI:** 10.1101/2022.07.22.501143

**Authors:** Danil V. Krivonos, Dmitry N. Konanov, Elena N. Ilina

## Abstract

Fungi are inseparable companions of human life, they can be found in both the environment and human organs including skin, respiratory tract and gut. Studies of fungal communities are of great interest to modern biology, partially due to their specific way of life and the presence of unique biochemical pathways they have. Fungi have been shown to be both producers of useful compounds, such as antibiotics and organic acids, and pathogens of various diseases. When considering the selected fungal community, in a number of cases it is rather difficult to evaluate its functional capabilities, which is partially caused by some technical difficulties in the analysis and annotation of whole eukaryotic genomes. In practice, the taxonomic composition of fungal communities is determined using short marker sequences. The most popular fungal taxonomy markers are ITS (internal transcribed spacer) sequences. Here, we present FunFun, the instrument that allows to evaluate the functional content of an individual fungus or mycobiome based on ITS sequencing data.

## 1. Introduction

Natural fungal communities have a lot of different biochemical pathways, including biosynthesis of antimicrobial substances, organic acids, and even toxins (Branco, 2019). At the same time, individual fungi from one community might drastically differ from each other in terms of their functional capabilities (Wisecaver et al., 2014).

Generally, while analyzing either individual fungi or their communities, researchers do not perform whole genome sequencing of eukaryotic cells and tend to infer their taxonomy using specific markers. This is caused by the high price of whole genome sequencing and some technical obstacles to work with eukaryotic cells. In practice, amplicon sequencing technologies are widely used for estimating fungal taxa composition. One of the most used taxonomy markers for fungi is ITS region (Blaalid et al., 2013; Mbareche et al., 2020; Schoch et al., 2012), which is useful in taxonomy inference but does not provide comprehensive information about genes content. However, in biotechnology it is essential to understand the biochemical capabilities of individual fungi, which might be applicable to industrial biotechnology (Kim et al., 2006). Information about gene content can be useful to solve ecological problems such as bioconversion of solid waste (Chilakamarry et al., 2022) or caffeine utilization (Zhou et al., 2018).

There are a number of tools designed to predict biochemical features of bacterial communities using 16S sequencing data (Douglas et al., 2020; Sun et al., 2020). At the same time, functional annotation of the fungal part of the community is usually performed using whole genome sequencing (WGS), since the number of precisely annotated fungi is not enough to build complex predictive models such as the machine learning algorithm implemented in PICRUST2 (Douglas et al., 2020). For this reason the fungal part of microbial communities often remains to be neglected.

Inspired by the idea of PICRUST2, we decided to develop an algorithm that is designed to solve a similar task for fungal datasets. The tool is based on the assumption that functional capabilities of fungi, unlike bacteria, are in strong correlation with their taxonomic affiliation which is according to published data describing individual fungal metabolic pathways (Slater and Birney, 2005). We implemented a modified *K* nearest neighbors (KNN) model and tested it using shuffle-split cross-validation and on ITS subset. The model has been implemented as a command-line tool called FunFun.

## 2. Methods

### 2.1 Data collection

To construct the model, we collected 9271 whole fungal genomes of different assembly levels from the GeneBank database. Next, we extracted ITS1-5.8S-ITS2 sequence fragments from the genomes via *in silico* PCR method with ITS1F: CTTGGTCATTTAGAGGAAGTAA and ITS4: TCCTCCGCTTATTGATATGC primers using the ipcress tool from the exonerate software package (Slater and Birney, 2005). After this stage, 6132 fungal genomes left, which might be the result of not successful primers annealing. Generally, *in silico* PCR does not guarantee the absolute correctness of extracted sequences. To validate ITS sequences we additionally used the ITSx software (Bengtsson-Palme et al., 2013) to ensure that the extracted fragments were ITS. Based on the ITSx annotations,, we built databases of individual ITS1 and ITS2 sequences, and full ITS1-5.8S-ITS2 fragments (they will be called Concatenates in the further text) belonging to 5882 fungal genomes.

The complete genome sequences were annotated using the Augustus tool (Keller et al., 2011) to ensure consistency of annotations. Predicted protein sequences were analyzed by the KofamKOALA tool (Aramaki et al., 2020) to get the functional annotations. Based on the results, we built gene content vectors constructed using the third level of the KoFAM hierarchy (430 vector components, presented in Supplementary 1). Each component in this vector represents the relative abundance of the corresponding KEGG orthology group. To avoid the appearance of non-fungal biochemical pathways, the KEGG orthology data were manually filtered by removing the 09160 Human Disease branch.

Additionally, we checked the quality of the gene content estimation based on Augustus genes annotations. 75 fungal assemblies for which curated protein sequences had been published were loaded from the RefSeq database. Here, to ensure the reliability of data, we considered only the protein sequence data obtained using the Eukaryotic Annotation Propagation Pipeline (“The NCBI Eukaryotic Genome Annotation Pipeline,” n.d.). Thereafter, for each assembly, we constructed two uniformic functional gene content profiles, where the first profile was based on curated protein sequences data from RefSeq and the second profile was built using Augustus gene annotations. Convergence of results was estimated using Pearson’s correlation coefficient for each assembly pair independently.

To preliminary validate our method of functional content profiling we used the t-SNE decomposition for gene content vectors. We expected that observed clusters would be associated with fungal taxonomy, which is according to literature data (Leroy et al., 2021; Wisecaver et al., 2014). To estimate the convergence of observed clusters and fungal taxonomy, the Rand Index (*RI*) was used.

### 2.2 Algorithm explanation

For each ITS sequence in the collected database, a relative abundance of each nucleotide k-mer is calculated (k was chosen to be 5). Thus, for each sequence we generate a k-mer relative abundances vector with length of 4^5^. In the further text we will name such k-mer abundances vectors as k-mers vectors. Thereafter, we evaluate cosine distance between the k-mers vector representing the target sequence and k-mers vectors for each fungal reference amplicon variant (RAV) from the database. Thus, we constructed a list of distances, which describe similarity between the target sequence and each RAV from the database.

To predict the fungal gene content, the algorithm searches for *K* nearest neighbors in the *ε* neighborhood using the generated distances list. Here, the *ε* neighborhood is the area, which limits the space where the neighbors searching will be performed. Number of neighbors not exceeding *K* and located in an area, limited by *ε* neighborhood, we call *K*_*chosen*_. Such an approach allows to prevent usage of too far sequences while gene content prediction. For each *i*-th fungi in our database we have a precalculated gene content profile 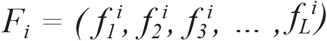, *F* is a vector with length *L* (*L* = 430). In *F*_*j*_, each 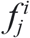 value is a fraction of individual j-th function while the sum of these values is equal 1. To calculate functional gene content profile for the target sample we average the value of each individual function among the *K*_*chosen*_ neighbors.

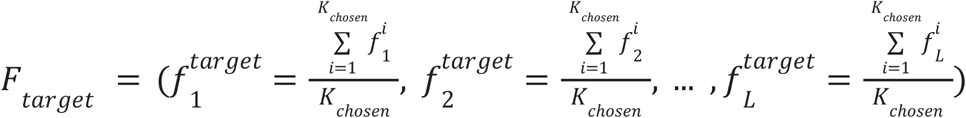, where *F*_*target*_ - predicted gene content profile of target fungi, *K*_*chosen*_ - number of neighbors not exceeding K and limited by ε neighborhood, 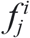-fraction of *j*-th function in the genome of *i-*th neighbored fungi from *K*_*chosen*_.

In cases, when the target sequence has neighbors with cosine distance to the target of 0 (that usually means the identity of ITS sequences), only these neighbors are selected for the prediction. is.

### 2.3 Validation

To test our algorithm we splitted our database on test/train data sets and tested using shuffle-split cross-validation. The train/test ratio was chosen to be 80/20, with 10 shuffling iterations. So, for each shuffling epoch 1176 testing samples were used. For each epoch, quality of prediction was estimated using a median *R*^*2*^ metric between real gene content vectors, which were built using whole genome annotations, and predicted vectors which were obtained with FunFun. The validation was performed for each amplicon type (ITS1, ITS2, Concatenate) independently.

## 3. Results

### 3.1. Software explanation

We have developed an algorithm aimed to predict fungal metabolic capabilities (gene content) using ITS sequencing data. As an input the algorithm can use ITS1, ITS2 or full size ITS clusters (concatenates). The algorithm has been realized as a command-line instrument which we named FunFun (Fungal Functional). As an input, the program takes the ITS sequence (or sequences) of the object (or objects) of interest in .fasta format and returns as an output the table, where the first column is the KEGG orthology group (which we call function), and the subsequent columns correspond to predicted gene content vector components for each sample. First of all, FunFun was developed to be capable of metagenomic analysis but can be useful in analysis of individual fungi as well.

Also, the tool has two hyperparameters *K* - maximum number of nearest neighbors selected and *ε*, defining the area where the *K* neighbors search is performed. In the case when the tool does not return a prediction for the analyzed sequence, the user can increase the ε value. However, it should be noted here that the greater the ε value the lower the expected prediction quality. On the other hand, as we observed the number of neighbors *K* does not drastically influence the prediction quality in general but in some cases might make the prediction more robust, especially while the target sequence has a lot of close neighbors. By default, *K* is set to be 10 and ε is set to be 0.5.

### 3.2. Validation results

The algorithm developed is based on the assumption that fungal metabolism correlates with their taxonomy, so, we expected that gene content profiles, which we calculated with our approach, also should be in strong correlation with fungal taxonomy. To verify this assumption, t-SNE decomposition of calculated gene content vectors and clustering with the HDBSCAN algorithm (Campello et al., 2015, 2013) were performed. The resulting clusters demonstrated a good convergence with taxonomy data (family level) (*RI* = 0.929). In part, the observed inaccuracy in clustering might be caused by some ambiguity of fungal taxonomy (Hawksworth, 2011).

We also checked the validity of Augustus annotations by comparing functional profiling of Augustus annotations with corresponding RefSeq protein data. The analysis showed a good value of Pearson correlation score (*r*=0.62). Thus, results of functional profiling obtained with Augustus are quite close to the results confirmed by experimental transcriptome data, so, we decided that Augustus is suitable as a genome annotator in our method.

Using the number of neighbors of *K* = 10, we tested our algorithm with a range of different *ε* values (Fig.3). At the validation stage we calculated the *R*^*2*^ value between the gene content vector, which we built on full genome data, and the gene content vector predicted on ITS sequence using FunFun. Also, we estimated the percentage of predicted samples from the test subset (functional gene content is predicted only if the target ITS has at least one neighbor in the ε neighborhood).

**FIGURE 1.**
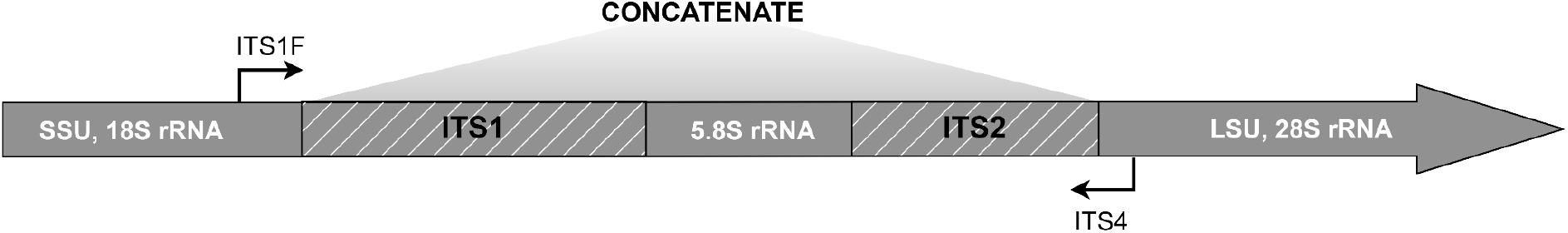
ITS cluster structure and primer scheme used.

**FIGURE 2.**
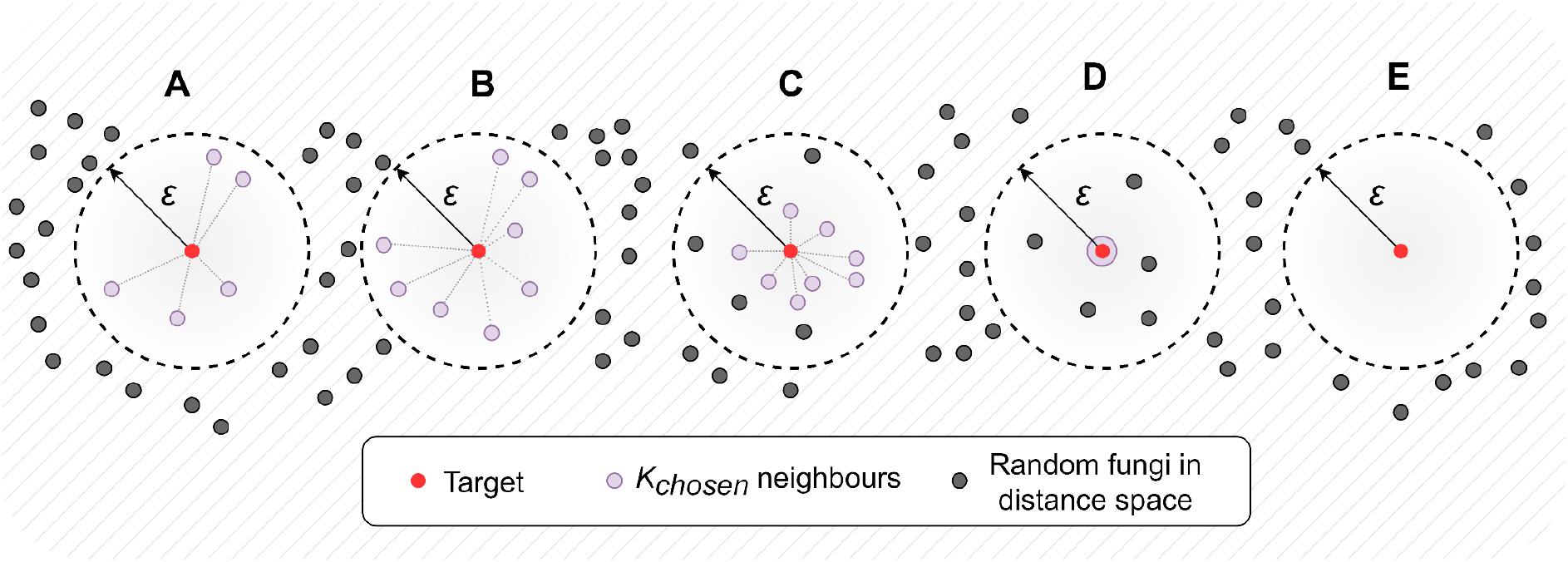
Possible cases of neighbors selection (in this example *K* = 8). A - when the number of reference amplicon variants (RAVs) included in the ε neighborhood is less than the specified number *K*; B - when the ε neighborhood includes exactly *K* neighbors; C - when the number of points included in the ε neighborhood is greater than *K*; D - when the target sequence has at least one RAV with identical k-mers vector; E - when there are no neighbors in the ε neighborhood.

**FIGURE 3.**
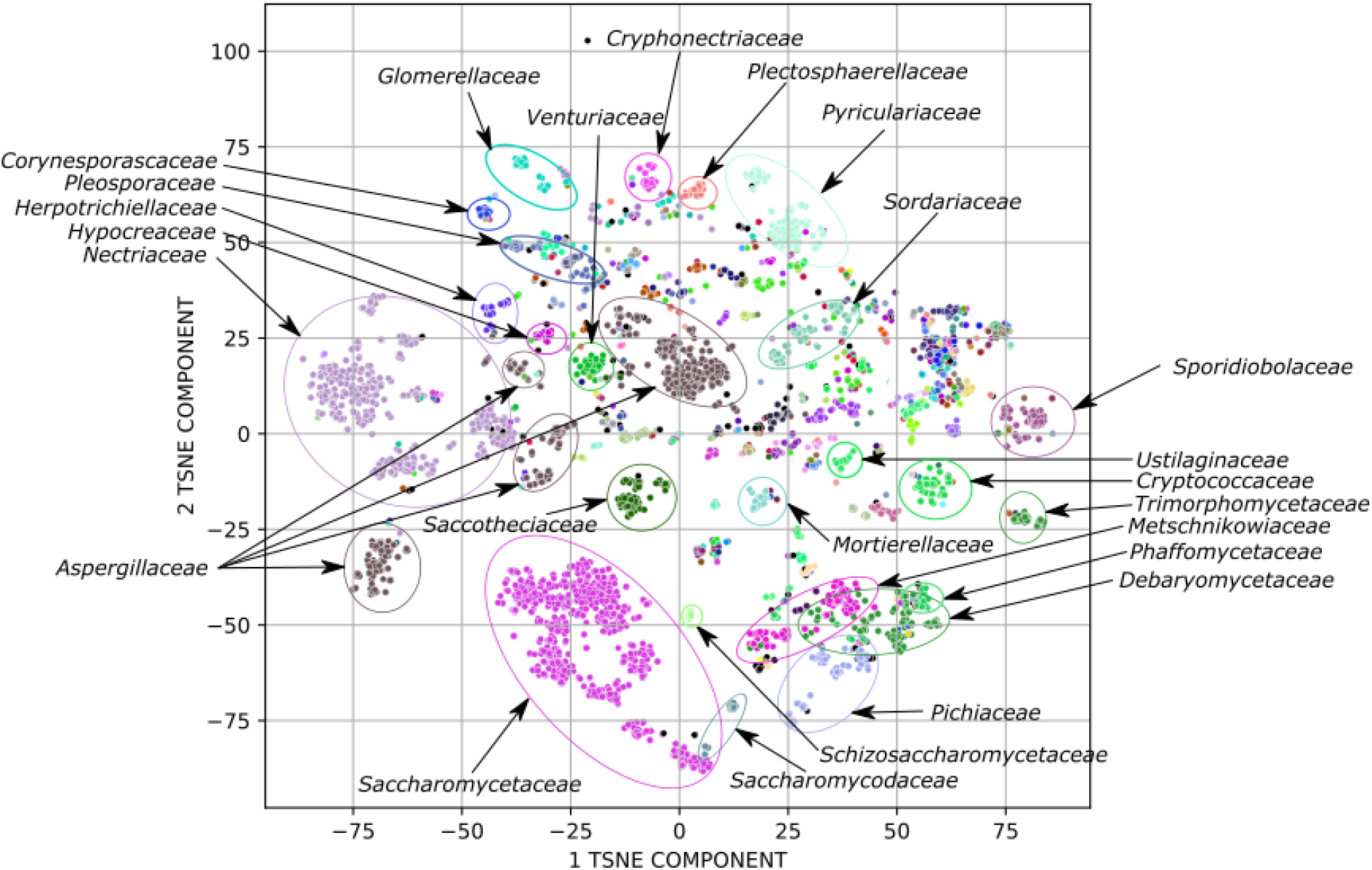
t-SNE decomposition of fungal functional profiles. Each color corresponds to an individual fungal taxon.

We found that it is more efficient to give the algorithm using as input data full size ITS clusters (concatenates). On average, the values of median *R*^*2*^ were at least 0.95 regardless of the *ε* value chosen. So, we set *ε* = 0.5 and *K* = 10 as default values. If necessary, the user is able to set custom values of them when running FunFun.

Quite high *R*^*2*^ values might be partially explained by conservative gene functions, which are common features of all fungi. Examples of these functions are glycolysis and citric cycle. So, we decided to additionally check how the method estimates relative abundances of the functions most variable among different fungal genomes. We chose such functions using the coefficient of variation, which is defined as the standard deviation normalized to the mean. We considered a function as highly variable if its coefficient of variation was greater than 10. As on the previous validation stage we calculated the median *R*^*2*^ value between real and predicted gene content vectors. In the result we obtained median *R*^*2*^ = 0.99 and percentage of predicted equals 98.6%.

Figure 5 illustrates convergence of real gene content profiles and corresponding profiles predicted with FunFun. For this assay we took a subset of 10 randomly chosen fungal organisms.

**FIGURE 4.**
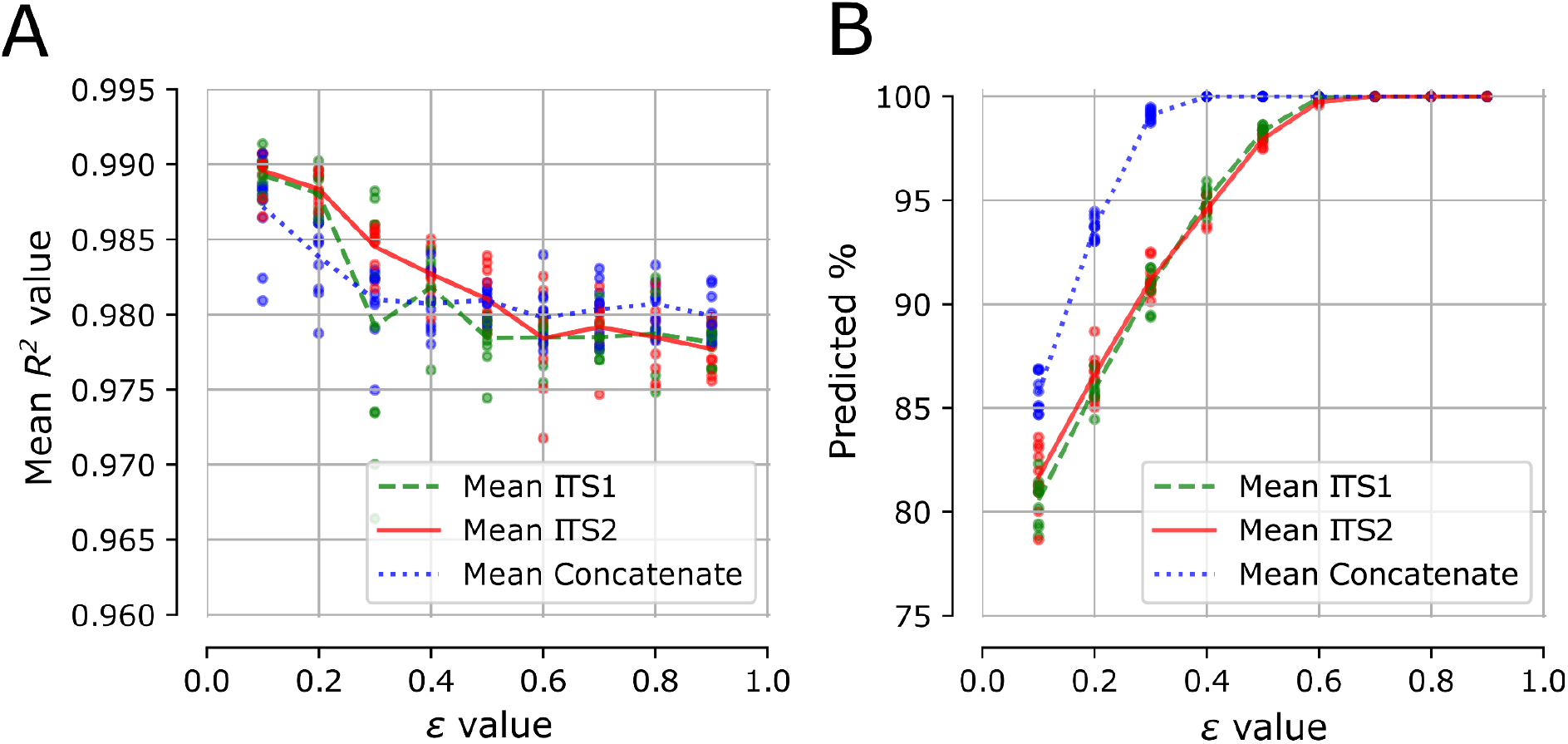
Validation results. **A**) Dependence of gene content prediction quality on the *ε* value. **B**) Dependence of the percentage of predictions on *ε* value (In some cases the *ε* neighborhood can include no one neighbors that does not allow to build predicted profiles with sufficient quality. Thus, the lower the *ε* value, the less the number of predicted profiles).

**FIGURE 5.**
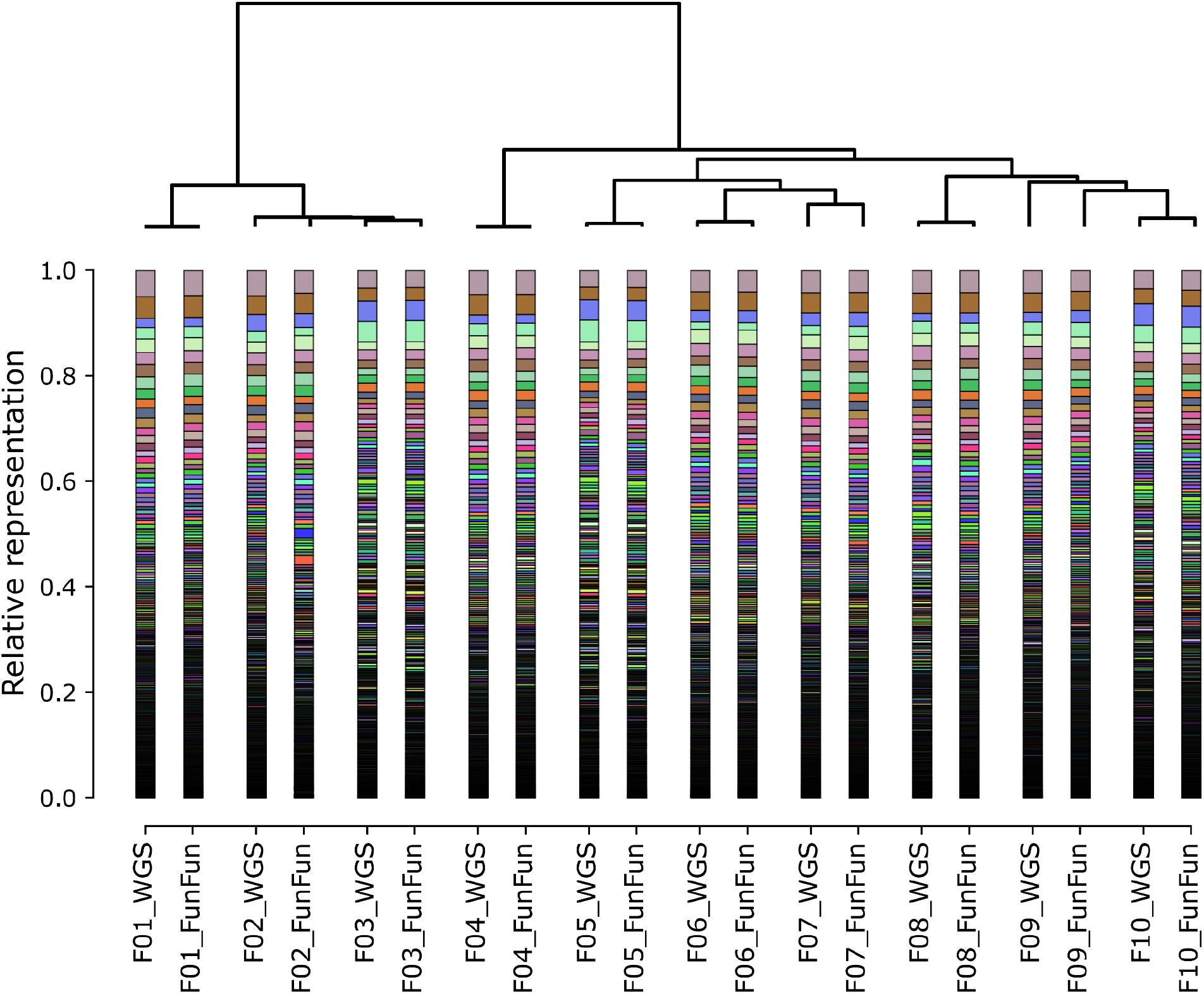
Comparison of gene content profiles obtained from whole genome sequence data and predicted on ITS data with FunFun. For each pair, the left bar represents the real relative abundances of corresponding functions in the genome while the right bar shows function fractions predicted with FunFun.

## Conclusion

We have developed FunFun, a novel tool designed for estimation of fungal functional content based on ITS amplicon sequencing data. It can be useful for estimation of fungal gene content for individual fungi as well as mycobioms. The tool can be installed via pip. Also,it can be downloaded from git https://github.com/DanilKrivonos/FunFun.

## CRediT authorship contribution statement

Danil V. Krivonos: Conceptualization, Software, Writing - original draft, Data curation, Formal analysis, Visualization, Dmitry N. Konanov: Conceptualization, Methodology, Formal analysis, Writing - original draft. Elena N. Ilina: Supervision, Writing - review & editing.

## References

Aramaki, T., Blanc-Mathieu, R., Endo, H., Ohkubo, K., Kanehisa, M., Goto, S., Ogata, H., 2020. KofamKOALA: KEGG Ortholog assignment based on profile HMM and adaptive score threshold. Bioinforma. Oxf. Engl. 36, 2251–2252. https://doi.org/10.1093/bioinformatics/btz859

Bengtsson-Palme, J., Ryberg, M., Hartmann, M., Branco, S., Wang, Z., Godhe, A., De Wit, P., Sánchez-García, M., Ebersberger, I., de Sousa, F., Amend, A., Jumpponen, A., Unterseher, M., Kristiansson, E., Abarenkov, K., Bertrand, Y.J.K., Sanli, K., Eriksson, K.M., Vik, U., Veldre, V., Nilsson, R.H., 2013. Improved software detection and extraction of ITS1 and ITS2 from ribosomal ITS sequences of fungi and other eukaryotes for analysis of environmental sequencing data. Methods Ecol. Evol. 4, 914–919. https://doi.org/10.1111/2041-210X.12073

Blaalid, R., Kumar, S., Nilsson, R.H., Abarenkov, K., Kirk, P.M., Kauserud, H., 2013. ITS1 versus ITS2 as DNA metabarcodes for fungi. Mol. Ecol. Resour. 13, 218–224. https://doi.org/10.1111/1755-0998.12065

Branco, S., 2019. Fungal diversity from communities to genes. Fungal Biol. Rev. 33, 225–237. https://doi.org/10.1016/j.fbr.2019.06.003

Campello, R.J.G.B., Moulavi, D., Sander, J., 2013. Density-Based Clustering Based on Hierarchical Density Estimates, in: Pei, J., Tseng, V.S., Cao, L., Motoda, H., Xu, G. (Eds.), Advances in Knowledge Discovery and Data Mining, Lecture Notes in Computer Science. Springer Berlin Heidelberg, Berlin, Heidelberg, pp. 160–172. https://doi.org/10.1007/978-3-642-37456-2_14

Campello, R.J.G.B., Moulavi, D., Zimek, A., Sander, J., 2015. Hierarchical Density Estimates for Data Clustering, Visualization, and Outlier Detection. ACM Trans. Knowl. Discov. Data 10, 5:1–5:51. https://doi.org/10.1145/2733381

Douglas, G.M., Maffei, V.J., Zaneveld, J.R., Yurgel, S.N., Brown, J.R., Taylor, C.M., Huttenhower, C., Langille, M.G.I., 2020. PICRUSt2 for prediction of metagenome functions. Nat. Biotechnol. 38, 685–688. https://doi.org/10.1038/s41587-020-0548-6

Hawksworth, D.L., 2011. A new dawn for the naming of fungi: impacts of decisions made in Melbourne in July 2011 on the future publication and regulation of fungal names1. IMA Fungus Glob. Mycol. J. 2, 155–162. https://doi.org/10.5598/imafungus.2011.02.02.06

Keller, O., Kollmar, M., Stanke, M., Waack, S., 2011. A novel hybrid gene prediction method employing protein multiple sequence alignments. Bioinforma. Oxf. Engl. 27, 757–763. https://doi.org/10.1093/bioinformatics/btr010

Kim, J.-W., Barrington, S., Sheppard, J., Lee, B., 2006. Nutrient optimization for the production of citric acid by Aspergillus niger NRRL 567 grown on peat moss enriched with glucose. Process Biochem. 41, 1253–1260. https://doi.org/10.1016/j.procbio.2005.12.021

Leroy, C., Maes, A.Q., Louisanna, E., Schimann, H., Séjalon-Delmas, N., 2021. Taxonomic, phylogenetic and functional diversity of root-associated fungi in bromeliads: effects of host identity, life forms and nutritional modes. New Phytol. 231, 1195–1209. https://doi.org/10.1111/nph.17288

Mbareche, H., Veillette, M., Bilodeau, G., Duchaine, C., 2020. Comparison of the performance of ITS1 and ITS2 as barcodes in amplicon-based sequencing of bioaerosols. PeerJ 8, e8523. https://doi.org/10.7717/peerj.8523

Schoch, C.L., Seifert, K.A., Huhndorf, S., Robert, V., Spouge, J.L., Levesque, C.A., Chen, W., Fungal Barcoding Consortium, Fungal Barcoding Consortium Author List, 2012. Nuclear ribosomal internal transcribed spacer (ITS) region as a universal DNA barcode marker for Fungi. Proc. Natl. Acad. Sci. U. S. A. 109, 6241–6246. https://doi.org/10.1073/pnas.1117018109

Slater, G.S.C., Birney, E., 2005. Automated generation of heuristics for biological sequence comparison. BMC Bioinformatics 6, 31. https://doi.org/10.1186/1471-2105-6-31

Sun, S., Jones, R.B., Fodor, A.A., 2020. Inference-based accuracy of metagenome prediction tools varies across sample types and functional categories. Microbiome 8, 46. https://doi.org/10.1186/s40168-020-00815-y

The NCBI Eukaryotic Genome Annotation Pipeline [WWW Document], n.d. URL https://www.ncbi.nlm.nih.gov/genome/annotation_euk/process/#references (accessed 7.18.22).

Wisecaver, J.H., Slot, J.C., Rokas, A., 2014. The evolution of fungal metabolic pathways. PLoS Genet. 10, e1004816. https://doi.org/10.1371/journal.pgen.1004816

Zhou, B., Ma, C., Wang, H., Xia, T., 2018. Biodegradation of caffeine by whole cells of tea-derived fungi Aspergillus sydowii, Aspergillus niger and optimization for caffeine degradation. BMC Microbiol. 18, 53. https://doi.org/10.1186/s12866-018-1194-8

