## Supplementary 1 for "FunFun: ITS-based functional annotator of fungal communities"

SUPPLEMENTARY 1 Gene contents classes (KEGG orthology group)

99984 Nucleotide metabolism

00601 Glycosphingolipid biosynthesis - lacto and neolacto series [PATH:ko00601]

03210 Viral fusion proteins [BR:ko03210]

00532 Glycosaminoglycan biosynthesis - chondroitin sulfate / dermatan sulfate [PATH:ko00532]

04610 Complement and coagulation cascades [PATH:ko04610]

00512 Mucin type O-glycan biosynthesis [PATH:ko00512]

04672 Intestinal immune network for IgA production [PATH:ko04672]

00905 Brassinosteroid biosynthesis [PATH:ko00905]

00942 Anthocyanin biosynthesis [PATH:ko00942]

00533 Glycosaminoglycan biosynthesis - keratan sulfate [PATH:ko00533]

00402 Benzoxazinoid biosynthesis [PATH:ko00402]

00943 Isoflavonoid biosynthesis [PATH:ko00943]

00196 Photosynthesis - antenna proteins [PATH:ko00196]

01056 Biosynthesis of type II polyketide backbone [PATH:ko01056]

00522 Biosynthesis of 12-, 14- and 16-membered macrolides [PATH:ko00522]

04052 Cytokines and growth factors [BR:ko04052]

03310 Nuclear receptors [BR:ko03310]

00365 Furfural degradation [PATH:ko00365]

00331 Clavulanic acid biosynthesis [PATH:ko00331]

00542 O-Antigen repeat unit biosynthesis [PATH:ko00542]

00571 Lipoarabinomannan (LAM) biosynthesis [PATH:ko00571]

04514 Cell adhesion molecules [PATH:ko04514]

00642 Ethylbenzene degradation [PATH:ko00642]

00572 Arabinogalactan biosynthesis - Mycobacterium [PATH:ko00572]

99979 Unclassified viral proteins

99986 Glycan metabolism

99998 Others

00195 Photosynthesis [PATH:ko00195]

00194 Photosynthesis proteins [BR:ko00194]

99993 Cell motility

04512 ECM-receptor interaction [PATH:ko04512]

04080 Neuroactive ligand-receptor interaction [PATH:ko04080]

00121 Secondary bile acid biosynthesis [PATH:ko00121]

02060 Phosphotransferase system (PTS) [PATH:ko02060]

00525 Acarbose and validamycin biosynthesis [PATH:ko00525]

00944 Flavone and flavonol biosynthesis [PATH:ko00944]

00997 Biosynthesis of various secondary metabolites - part 3 [PATH:ko00997]

02040 Flagellar assembly [PATH:ko02040]

04640 Hematopoietic cell lineage [PATH:ko04640]

01055 Biosynthesis of vancomycin group antibiotics [PATH:ko01055]

04030 G protein-coupled receptors [BR:ko04030]

00253 Tetracycline biosynthesis [PATH:ko00253]

00902 Monoterpenoid biosynthesis [PATH:ko00902]

00540 Lipopolysaccharide biosynthesis [PATH:ko00540]

01005 Lipopolysaccharide biosynthesis proteins [BR:ko01005]

01052 Type I polyketide structures [PATH:ko01052]

02030 Bacterial chemotaxis [PATH:ko02030]

09113 Global maps only

00534 Glycosaminoglycan biosynthesis - heparan sulfate / heparin [PATH:ko00534]

99974 Translation

00473 D-Alanine metabolism [PATH:ko00473]

00633 Nitrotoluene degradation [PATH:ko00633]

02035 Bacterial motility proteins [BR:ko02035]

00550 Peptidoglycan biosynthesis [PATH:ko00550]

00471 D-Glutamine and D-glutamate metabolism [PATH:ko00471]

01057 Biosynthesis of type II polyketide products [PATH:ko01057]

01051 Biosynthesis of ansamycins [PATH:ko01051]

05111 Biofilm formation - Vibrio cholerae [PATH:ko05111]

00523 Polyketide sugar unit biosynthesis [PATH:ko00523]

00945 Stilbenoid, diarylheptanoid and gingerol biosynthesis [PATH:ko00945]

99992 Structural proteins

04075 Plant hormone signal transduction [PATH:ko04075]

00332 Carbapenem biosynthesis [PATH:ko00332]

00908 Zeatin biosynthesis [PATH:ko00908]

04054 Pattern recognition receptors [BR:ko04054]

00984 Steroid degradation [PATH:ko00984]

00941 Flavonoid biosynthesis [PATH:ko00941]

00073 Cutin, suberine and wax biosynthesis [PATH:ko00073]

00515 Mannose type O-glycan biosynthesis [PATH:ko00515]

04711 Circadian rhythm - fly [PATH:ko04711]

02042 Bacterial toxins [BR:ko02042]

00622 Xylene degradation [PATH:ko00622]

04744 Phototransduction [PATH:ko04744]

04742 Taste transduction [PATH:ko04742]

99999 Others

00232 Caffeine metabolism [PATH:ko00232]

00535 Proteoglycans [BR:ko00535]

04215 Apoptosis - multiple species [PATH:ko04215]

01054 Nonribosomal peptide structures [PATH:ko01054]

04927 Cortisol synthesis and secretion [PATH:ko04927]

99975 Protein processing

99976 Replication and repair

04620 Toll-like receptor signaling pathway [PATH:ko04620]

99985 Amino acid metabolism

04964 Proximal tubule bicarbonate reclamation [PATH:ko04964]

03070 Bacterial secretion system [PATH:ko03070]

00536 Glycosaminoglycan binding proteins [BR:ko00536]

01053 Biosynthesis of siderophore group nonribosomal peptides [PATH:ko01053]

99981 Carbohydrate metabolism

00785 Lipoic acid metabolism [PATH:ko00785]

00604 Glycosphingolipid biosynthesis - ganglio series [PATH:ko00604]

01011 Peptidoglycan biosynthesis and degradation proteins [BR:ko01011]

04913 Ovarian steroidogenesis [PATH:ko04913]

04112 Cell cycle - Caulobacter [PATH:ko04112]

04712 Circadian rhythm - plant [PATH:ko04712]

00405 Phenazine biosynthesis [PATH:ko00405]

03230 Viral genome structure [PATH:ko03230]

00966 Glucosinolate biosynthesis [PATH:ko00966]

00364 Fluorobenzoate degradation [PATH:ko00364]

00140 Steroid hormone biosynthesis [PATH:ko00140]

00120 Primary bile acid biosynthesis [PATH:ko00120]

99994 Others

00403 Indole diterpene alkaloid biosynthesis [PATH:ko00403]

02026 Biofilm formation - Escherichia coli [PATH:ko02026]

03200 Viral proteins [BR:ko03200]

04515 Cell adhesion molecules [BR:ko04515]

02048 Prokaryotic defense system [BR:ko02048]

00281 Geraniol degradation [PATH:ko00281]

99973 Transcription

04740 Olfactory transduction [PATH:ko04740]

99987 Cofactor metabolism

99978 Cell growth

00363 Bisphenol degradation [PATH:ko00363]

04745 Phototransduction - fly [PATH:ko04745]

02025 Biofilm formation - Pseudomonas aeruginosa [PATH:ko02025]

00999 Biosynthesis of various secondary metabolites - part 1 [PATH:ko00999]

00440 Phosphonate and phosphinate metabolism [PATH:ko00440]

00472 D-Arginine and D-ornithine metabolism [PATH:ko00472]

00906 Carotenoid biosynthesis [PATH:ko00906]

01059 Biosynthesis of enediyne antibiotics [PATH:ko01059]

00603 Glycosphingolipid biosynthesis - globo and isoglobo series [PATH:ko00603]

00909 Sesquiterpenoid and triterpenoid biosynthesis [PATH:ko00909]

04668 TNF signaling pathway [PATH:ko04668]

01504 Antimicrobial resistance genes [BR:ko01504]

99982 Energy metabolism

04929 GnRH secretion [PATH:ko04929]

00404 Staurosporine biosynthesis [PATH:ko00404]

04614 Renin-angiotensin system [PATH:ko04614]

00311 Penicillin and cephalosporin biosynthesis [PATH:ko00311]

04658 Th1 and Th2 cell differentiation [PATH:ko04658]

00791 Atrazine degradation [PATH:ko00791]

04911 Insulin secretion [PATH:ko04911]

00514 Other types of O-glycan biosynthesis [PATH:ko00514]

04064 NF-kappa B signaling pathway [PATH:ko04064]

00901 Indole alkaloid biosynthesis [PATH:ko00901]

00524 Neomycin, kanamycin and gentamicin biosynthesis [PATH:ko00524]

04320 Dorso-ventral axis formation [PATH:ko04320]

04923 Regulation of lipolysis in adipocytes [PATH:ko04923]

99995 Signaling proteins

04380 Osteoclast differentiation [PATH:ko04380]

04624 Toll and Imd signaling pathway [PATH:ko04624]

00062 Fatty acid elongation [PATH:ko00062]

99983 Lipid metabolism

00591 Linoleic acid metabolism [PATH:ko00591]

00261 Monobactam biosynthesis [PATH:ko00261]

04330 Notch signaling pathway [PATH:ko04330]

99977 Transport

00965 Betalain biosynthesis [PATH:ko00965]

04550 Signaling pathways regulating pluripotency of stem cells [PATH:ko04550]

04960 Aldosterone-regulated sodium reabsorption [PATH:ko04960]

04979 Cholesterol metabolism [PATH:ko04979]

00531 Glycosaminoglycan degradation [PATH:ko00531]

04091 Lectins [BR:ko04091]

99996 General function prediction only

00904 Diterpenoid biosynthesis [PATH:ko00904]

04924 Renin secretion [PATH:ko04924]

04350 TGF-beta signaling pathway [PATH:ko04350]

04971 Gastric acid secretion [PATH:ko04971]

04975 Fat digestion and absorption [PATH:ko04975]

04630 JAK-STAT signaling pathway [PATH:ko04630]

00401 Novobiocin biosynthesis [PATH:ko00401]

04391 Hippo signaling pathway - fly [PATH:ko04391]

04713 Circadian entrainment [PATH:ko04713]

04520 Adherens junction [PATH:ko04520]

04122 Sulfur relay system [PATH:ko04122]

00998 Biosynthesis of various secondary metabolites - part 2 [PATH:ko00998]

04622 RIG-I-like receptor signaling pathway [PATH:ko04622]

04626 Plant-pathogen interaction [PATH:ko04626]

04710 Circadian rhythm [PATH:ko04710]

00430 Taurine and hypotaurine metabolism [PATH:ko00430]

00750 Vitamin B6 metabolism [PATH:ko00750]

04978 Mineral absorption [PATH:ko04978]

04917 Prolactin signaling pathway [PATH:ko04917]

04040 Ion channels [BR:ko04040]

04659 Th17 cell differentiation [PATH:ko04659]

00361 Chlorocyclohexane and chlorobenzene degradation [PATH:ko00361]

04730 Long-term depression [PATH:ko04730]

04750 Inflammatory mediator regulation of TRP channels [PATH:ko04750]

04662 B cell receptor signaling pathway [PATH:ko04662]

00660 C5-Branched dibasic acid metabolism [PATH:ko00660]

04670 Leukocyte transendothelial migration [PATH:ko04670]

04974 Protein digestion and absorption [PATH:ko04974]

04623 Cytosolic DNA-sensing pathway [PATH:ko04623]

04392 Hippo signaling pathway - multiple species [PATH:ko04392]

04918 Thyroid hormone synthesis [PATH:ko04918]

04340 Hedgehog signaling pathway [PATH:ko04340]

00541 O-Antigen nucleotide sugar biosynthesis [PATH:ko00541]

00623 Toluene degradation [PATH:ko00623]

04341 Hedgehog signaling pathway - fly [PATH:ko04341]

00592 alpha-Linolenic acid metabolism [PATH:ko00592]

04370 VEGF signaling pathway [PATH:ko04370]

00930 Caprolactam degradation [PATH:ko00930]

01040 Biosynthesis of unsaturated fatty acids [PATH:ko01040]

04973 Carbohydrate digestion and absorption [PATH:ko04973]

04970 Salivary secretion [PATH:ko04970]

04611 Platelet activation [PATH:ko04611]

04664 Fc epsilon RI signaling pathway [PATH:ko04664]

00521 Streptomycin biosynthesis [PATH:ko00521]

00643 Styrene degradation [PATH:ko00643]

04015 Rap1 signaling pathway [PATH:ko04015]

04130 SNARE interactions in vesicular transport [PATH:ko04130]

04216 Ferroptosis [PATH:ko04216]

04650 Natural killer cell mediated cytotoxicity [PATH:ko04650]

04260 Cardiac muscle contraction [PATH:ko04260]

04720 Long-term potentiation [PATH:ko04720]

04657 IL-17 signaling pathway [PATH:ko04657]

04920 Adipocytokine signaling pathway [PATH:ko04920]

04966 Collecting duct acid secretion [PATH:ko04966]

04928 Parathyroid hormone synthesis, secretion and action [PATH:ko04928]

00590 Arachidonic acid metabolism [PATH:ko00590]

04625 C-type lectin receptor signaling pathway [PATH:ko04625]

03450 Non-homologous end-joining [PATH:ko03450]

00981 Insect hormone biosynthesis [PATH:ko00981]

04925 Aldosterone synthesis and secretion [PATH:ko04925]

00740 Riboflavin metabolism [PATH:ko00740]

03060 Protein export [PATH:ko03060]

04016 MAPK signaling pathway - plant [PATH:ko04016]

04270 Vascular smooth muscle contraction [PATH:ko04270]

04926 Relaxin signaling pathway [PATH:ko04926]

04916 Melanogenesis [PATH:ko04916]

04972 Pancreatic secretion [PATH:ko04972]

00624 Polycyclic aromatic hydrocarbon degradation [PATH:ko00624]

00621 Dioxin degradation [PATH:ko00621]

04390 Hippo signaling pathway [PATH:ko04390]

04012 ErbB signaling pathway [PATH:ko04012]

00511 Other glycan degradation [PATH:ko00511]

00720 Carbon fixation pathways in prokaryotes [PATH:ko00720]

04961 Endocrine and other factor-regulated calcium reabsorption [PATH:ko04961]

04210 Apoptosis [PATH:ko04210]

00565 Ether lipid metabolism [PATH:ko00565]

04726 Serotonergic synapse [PATH:ko04726]

04725 Cholinergic synapse [PATH:ko04725]

04115 p53 signaling pathway [PATH:ko04115]

00960 Tropane, piperidine and pyridine alkaloid biosynthesis [PATH:ko00960]

00670 One carbon pool by folate [PATH:ko00670]

04214 Apoptosis - fly [PATH:ko04214]

02044 Secretion system [BR:ko02044]

04612 Antigen processing and presentation [PATH:ko04612]

00903 Limonene and pinene degradation [PATH:ko00903]

00730 Thiamine metabolism [PATH:ko00730]

01006 Prenyltransferases [BR:ko01006]

04660 T cell receptor signaling pathway [PATH:ko04660]

04360 Axon guidance [PATH:ko04360]

00450 Selenocompound metabolism [PATH:ko00450]

04935 Growth hormone synthesis, secretion and action [PATH:ko04935]

99988 Secondary metabolism

04022 cGMP-PKG signaling pathway [PATH:ko04022]

04013 MAPK signaling pathway - fly [PATH:ko04013]

00563 Glycosylphosphatidylinositol (GPI)-anchor biosynthesis [PATH:ko00563]

04962 Vasopressin-regulated water reabsorption [PATH:ko04962]

04727 GABAergic synapse [PATH:ko04727]

04062 Chemokine signaling pathway [PATH:ko04062]

04724 Glutamatergic synapse [PATH:ko04724]

04977 Vitamin digestion and absorption [PATH:ko04977]

04261 Adrenergic signaling in cardiomyocytes [PATH:ko04261]

04137 Mitophagy - animal [PATH:ko04137]

04510 Focal adhesion [PATH:ko04510]

04540 Gap junction [PATH:ko04540]

04090 CD molecules [BR:ko04090]

00333 Prodigiosin biosynthesis [PATH:ko00333]

04915 Estrogen signaling pathway [PATH:ko04915]

00600 Sphingolipid metabolism [PATH:ko00600]

00900 Terpenoid backbone biosynthesis [PATH:ko00900]

04912 GnRH signaling pathway [PATH:ko04912]

00254 Aflatoxin biosynthesis [PATH:ko00254]

00983 Drug metabolism - other enzymes [PATH:ko00983]

00980 Metabolism of xenobiotics by cytochrome P450 [PATH:ko00980]

04310 Wnt signaling pathway [PATH:ko04310]

04020 Calcium signaling pathway [PATH:ko04020]

02022 Two-component system [BR:ko02022]

04722 Neurotrophin signaling pathway [PATH:ko04722]

04976 Bile secretion [PATH:ko04976]

04371 Apelin signaling pathway [PATH:ko04371]

00790 Folate biosynthesis [PATH:ko00790]

04024 cAMP signaling pathway [PATH:ko04024]

00982 Drug metabolism - cytochrome P450 [PATH:ko00982]

00290 Valine, leucine and isoleucine biosynthesis [PATH:ko00290]

04621 NOD-like receptor signaling pathway [PATH:ko04621]

00100 Steroid biosynthesis [PATH:ko00100]

04666 Fc gamma R-mediated phagocytosis [PATH:ko04666]

00510 N-Glycan biosynthesis [PATH:ko00510]

00950 Isoquinoline alkaloid biosynthesis [PATH:ko00950]

00910 Nitrogen metabolism [PATH:ko00910]

02024 Quorum sensing [PATH:ko02024]

00130 Ubiquinone and other terpenoid-quinone biosynthesis [PATH:ko00130]

03020 RNA polymerase [PATH:ko03020]

00860 Porphyrin and chlorophyll metabolism [PATH:ko00860]

04072 Phospholipase D signaling pathway [PATH:ko04072]

00780 Biotin metabolism [PATH:ko00780]

03410 Base excision repair [PATH:ko03410]

03320 PPAR signaling pathway [PATH:ko03320]

00626 Naphthalene degradation [PATH:ko00626]

00300 Lysine biosynthesis [PATH:ko00300]

00710 Carbon fixation in photosynthetic organisms [PATH:ko00710]

04728 Dopaminergic synapse [PATH:ko04728]

04031 GTP-binding proteins [BR:ko04031]

04922 Glucagon signaling pathway [PATH:ko04922]

04361 Axon regeneration [PATH:ko04361]

03440 Homologous recombination [PATH:ko03440]

04066 HIF-1 signaling pathway [PATH:ko04066]

04217 Necroptosis [PATH:ko04217]

04721 Synaptic vesicle cycle [PATH:ko04721]

00513 Various types of N-glycan biosynthesis [PATH:ko00513]

04921 Oxytocin signaling pathway [PATH:ko04921]

04914 Progesterone-mediated oocyte maturation [PATH:ko04914]

04530 Tight junction [PATH:ko04530]

04136 Autophagy - other [PATH:ko04136]

00920 Sulfur metabolism [PATH:ko00920]

04068 FoxO signaling pathway [PATH:ko04068]

04613 Neutrophil extracellular trap formation [PATH:ko04613]

00830 Retinol metabolism [PATH:ko00830]

00537 Glycosylphosphatidylinositol (GPI)-anchored proteins [BR:ko00537]

03022 Basal transcription factors [PATH:ko03022]

99997 Function unknown

00053 Ascorbate and aldarate metabolism [PATH:ko00053]

04211 Longevity regulating pathway [PATH:ko04211]

04014 Ras signaling pathway [PATH:ko04014]

04212 Longevity regulating pathway - worm [PATH:ko04212]

04010 MAPK signaling pathway [PATH:ko04010]

03430 Mismatch repair [PATH:ko03430]

00362 Benzoate degradation [PATH:ko00362]

04810 Regulation of actin cytoskeleton [PATH:ko04810]

00760 Nicotinate and nicotinamide metabolism [PATH:ko00760]

00340 Histidine metabolism [PATH:ko00340]

03460 Fanconi anemia pathway [PATH:ko03460]

00625 Chloroalkane and chloroalkene degradation [PATH:ko00625]

00040 Pentose and glucuronate interconversions [PATH:ko00040]

04990 Domain-containing proteins not elsewhere classified [BR:ko04990]

00460 Cyanoamino acid metabolism [PATH:ko00460]

04070 Phosphatidylinositol signaling system [PATH:ko04070]

00030 Pentose phosphate pathway [PATH:ko00030]

00400 Phenylalanine, tyrosine and tryptophan biosynthesis [PATH:ko00400]

00220 Arginine biosynthesis [PATH:ko00220]

04723 Retrograde endocannabinoid signaling [PATH:ko04723]

00020 Citrate cycle (TCA cycle) [PATH:ko00020]

00480 Glutathione metabolism [PATH:ko00480]

00940 Phenylpropanoid biosynthesis [PATH:ko00940]

04152 AMPK signaling pathway [PATH:ko04152]

00680 Methane metabolism [PATH:ko00680]

04218 Cellular senescence [PATH:ko04218]

00240 Pyrimidine metabolism [PATH:ko00240]

04919 Thyroid hormone signaling pathway [PATH:ko04919]

00061 Fatty acid biosynthesis [PATH:ko00061]

00770 Pantothenate and CoA biosynthesis [PATH:ko00770]

00052 Galactose metabolism [PATH:ko00052]

00360 Phenylalanine metabolism [PATH:ko00360]

04145 Phagosome [PATH:ko04145]

02020 Two-component system [PATH:ko02020]

00410 beta-Alanine metabolism [PATH:ko00410]

01008 Polyketide biosynthesis proteins [BR:ko01008]

00640 Propanoate metabolism [PATH:ko00640]

00627 Aminobenzoate degradation [PATH:ko00627]

04213 Longevity regulating pathway - multiple species [PATH:ko04213]

04151 PI3K-Akt signaling pathway [PATH:ko04151]

00561 Glycerolipid metabolism [PATH:ko00561]

04114 Oocyte meiosis [PATH:ko04114]

04139 Mitophagy - yeast [PATH:ko04139]

04910 Insulin signaling pathway [PATH:ko04910]

00199 Cytochrome P450 [BR:ko00199]

03050 Proteasome [PATH:ko03050]

00051 Fructose and mannose metabolism [PATH:ko00051]

04140 Autophagy - animal [PATH:ko04140]

04071 Sphingolipid signaling pathway [PATH:ko04071]

03030 DNA replication [PATH:ko03030]

03420 Nucleotide excision repair [PATH:ko03420]

00562 Inositol phosphate metabolism [PATH:ko00562]

00071 Fatty acid degradation [PATH:ko00071]

00310 Lysine degradation [PATH:ko00310]

00564 Glycerophospholipid metabolism [PATH:ko00564]

00650 Butanoate metabolism [PATH:ko00650]

00350 Tyrosine metabolism [PATH:ko00350]

01004 Lipid biosynthesis proteins [BR:ko01004]

00630 Glyoxylate and dicarboxylate metabolism [PATH:ko00630]

00250 Alanine, aspartate and glutamate metabolism [PATH:ko00250]

04142 Lysosome [PATH:ko04142]

03015 mRNA surveillance pathway [PATH:ko03015]

00280 Valine, leucine and isoleucine degradation [PATH:ko00280]

03018 RNA degradation [PATH:ko03018]

04150 mTOR signaling pathway [PATH:ko04150]

00190 Oxidative phosphorylation [PATH:ko00190]

00970 Aminoacyl-tRNA biosynthesis [PATH:ko00970]

02010 ABC transporters [PATH:ko02010]

03010 Ribosome [PATH:ko03010]

03051 Proteasome [BR:ko03051]

00380 Tryptophan metabolism [PATH:ko00380]

04120 Ubiquitin mediated proteolysis [PATH:ko04120]

00330 Arginine and proline metabolism [PATH:ko00330]

03011 Ribosome [BR:ko03011]

01003 Glycosyltransferases [BR:ko01003]

00270 Cysteine and methionine metabolism [PATH:ko00270]

04110 Cell cycle [PATH:ko04110]

04146 Peroxisome [PATH:ko04146]

00520 Amino sugar and nucleotide sugar metabolism [PATH:ko00520]

00010 Glycolysis / Gluconeogenesis [PATH:ko00010]

00230 Purine metabolism [PATH:ko00230]

04714 Thermogenesis [PATH:ko04714]

00500 Starch and sucrose metabolism [PATH:ko00500]

03008 Ribosome biogenesis in eukaryotes [PATH:ko03008]

03013 Nucleocytoplasmic transport [PATH:ko03013]

00260 Glycine, serine and threonine metabolism [PATH:ko00260]

03012 Translation factors [BR:ko03012]

04144 Endocytosis [PATH:ko04144]

01007 Amino acid related enzymes [BR:ko01007]

00620 Pyruvate metabolism [PATH:ko00620]

04138 Autophagy - yeast [PATH:ko04138]

04141 Protein processing in endoplasmic reticulum [PATH:ko04141]

04812 Cytoskeleton proteins [BR:ko04812]

04113 Meiosis - yeast [PATH:ko04113]

03040 Spliceosome [PATH:ko03040]

04011 MAPK signaling pathway - yeast [PATH:ko04011]

03000 Transcription factors [BR:ko03000]

03032 DNA replication proteins [BR:ko03032]

04111 Cell cycle - yeast [PATH:ko04111]

01009 Protein phosphatases and associated proteins [BR:ko01009]

03110 Chaperones and folding catalysts [BR:ko03110]

01001 Protein kinases [BR:ko01001]

03021 Transcription machinery [BR:ko03021]

03016 Transfer RNA biogenesis [BR:ko03016]

03041 Spliceosome [BR:ko03041]

04121 Ubiquitin system [BR:ko04121]

03029 Mitochondrial biogenesis [BR:ko03029]

01002 Peptidases and inhibitors [BR:ko01002]

03019 Messenger RNA biogenesis [BR:ko03019]

03400 DNA repair and recombination proteins [BR:ko03400]

03009 Ribosome biogenesis [BR:ko03009]

04147 Exosome [BR:ko04147]

99980 Enzymes with EC numbers

02000 Transporters [BR:ko02000]

03036 Chromosome and associated proteins [BR:ko03036]

04131 Membrane trafficking [BR:ko04131]
